# Inferring the demographic history of Chinese and Indian rhesus macaque (*Macaca mulatta*) populations from PacBio HiFi long-read sequencing data

**DOI:** 10.64898/2026.05.25.727731

**Authors:** Erangi J. Heenkenda, Cyril J. Versoza, John W. Terbot, Vivak Soni, Gabriella J. Spatola, Susanne P. Pfeifer, Jeffrey D. Jensen

## Abstract

The rhesus macaque (*Macaca mulatta*) is one of the most widely used animal models in biomedical research, both as it resembles humans in key biological aspects and as it is characterized by a broad geographic range. Most of the individuals housed in U.S. research colonies have been sampled from either China or India, though notably the source population of these animals has significantly shifted over time. Given the substantial genetic and immunological differences between these populations, a deeper understanding of the underlying population structure is critically important for biomedical interpretation. Despite this, the demographic histories of these two populations remain poorly resolved. Here, we present an analysis of whole-genome, PacBio HiFi long-read sequencing data from ten unrelated individuals of each population, applying four related model- and non-model based demographic inference approaches, in order to reconstruct their ancestral history. We evaluated the fit of the subsequently estimated models against the empirical data, and incorporated underlying uncertainty in the mutation rates used for scaling. We inferred a well-fitting population history characterized by substantial structure between Chinese and Indian populations, with a split time ∼140,000 generations ago from an ancestral population of ∼65,000 individuals. We additionally inferred the subsequent history of size change within, and gene flow between, these populations, reaching the current estimated sizes of ∼220,000 individuals in the Chinese population and ∼14,000 individuals in the Indian population. The robust baseline demographic model established in this study will serve as a valuable resource for future research on this species, including for improved fine-scale recombination mapping, selection inference, and association studies.

## INTRODUCTION

Rhesus macaques (*Macaca mulatta*) — cercopithecoids with the largest natural geographic range of any non-human primate — are found from western India and Pakistan to the Pacific coast of China, and south into Vietnam and Thailand (Groves 2001). Previous inference has suggested a common ancestor with humans roughly 25–35 million years ago (Kumar and Hedges 1998; Perelman et al. 2011; Chintalapati and Moorjani 2020), and they have long been noted in the biomedical literature for their similarity to humans in key physiological and neurological traits as well as in their susceptibility to infectious and metabolic diseases (Rogers 2022). Owing to this resemblance with humans and widespread distribution, rhesus macaques are among the most extensively studied and utilized primate species in research (Rogers 2022). For example, they continue to serve as primary animal models for addressing fundamental questions in developmental psychology and social behavior, and are widely used in studies of infectious diseases, including HIV-AIDS (Liang et al. 2019), tuberculosis (Sharan et al. 2020), and more recently SARS-CoV-2 and COVID-19 (Klasse et al. 2021).

Despite this focus, numerous open questions remain pertaining to the underlying population structure and demographic dynamics of the species (Zhou et al. 2024; Terbot et al. 2025b). For example, the nature and degree of population subdivision across the rhesus macaque range continues to be the subject of debate. Early studies based on morphological measurements and pelage coloration proposed the existence of multiple subspecies, with suggestions of up to six putative subspecies in mainland China alone (Jiang et al. 1991). Subsequent work based on both mitochondrial (Zhang and Shi 1993) and nuclear (Liu et al. 2018) DNA largely supported this subdivision; however, more recent work opposes this claim, with results instead suggesting a single, interbreeding population within China, characterized by a population size change history closely linked to historical glaciation patterns (Terbot et al. 2025b). In contrast, comparisons between Indian- and Chinese-origin rhesus macaques have consistently revealed significant genetic divergence, with morphometric and genetic evidence supporting the existence of two distinct populations (Clarke and O’Neil 1999; Smith and McDonough 2005; Ferguson et al. 2007; Zhou et al. 2024). Specifically, previous genomic studies have suggested minimal gene flow between Chinese and Indian populations over the past 0.16 million years (Hernandez et al. 2007), and a mean *F_ST_* value of 0.14 has been reported between the two groups (Cooper et al. 2022). This population structuring has also been supported by recent work identifying numerous population-specific structural variants (Maruki et al. 2026). Notably, this structuring also has important practical implications for biomedical studies, as the origins of the individuals commonly used in research have shifted over time. Specifically, India, which had been the primary source of the populations utilized in research in the United States, suspended exports in 1978. After this time research facilities began importing more heavily from China (Cooper et al. 2022), thus altering the composition of the colonies, which to date are largely maintained through domestic breeding.

Key behavioral and phenotypic differences have been described between these populations, with comparative research on individuals of Indian- and Chinese-origin having expanded along with the accompanying growth in genomic resources over the past two decades. These studies have sought to characterize the extent to which functional genetic variation impacts physiology, immunology, and behavior (Cooper et al. 2022), with a particular focus on population-specific differences in disease pathogenesis, blood chemistry, the major histocompatibility complex, as well as general aspects of behavior and temperament (Champoux et al. 1994, 1996; Binhua et al. 2002; Ma et al. 2009; Jiang et al. 2013). At the same time, comparatively little research has been done to infer the underlying demographic dynamics characterizing the Indian and Chinese populations, despite the importance of this knowledge for the biological interpretation of genome-wide association and genotype-to-phenotype studies. Early work based upon 1,476 single nucleotide polymorphisms (SNPs) suggested a divergence time of 162,000 years (Hernandez et al. 2007), while recent whole-genome sequencing and alternative inference frameworks have reached broadly similar conclusions but with considerable uncertainty (Xue et al. 2016; Zhou et al. 2024). More generally, these existing studies differ in the markers, models, and analytical approaches used, resulting in an incomplete or inconsistent understanding of the history of the two populations, and have yet to propose a well-fitting demographic history encompassing population splits, size changes, and historical gene flow.

In order to better illuminate the demographic dynamics of this widely-studied species, we have utilized newly generated, whole-genome, high-fidelity, long-read sequencing data from 20 unrelated individuals (ten Chinese-origin and ten Indian-origin) to firstly quantify patterns of genetic diversity at putatively neutral, intergenic regions of the genome. Using this data, we evaluated levels of population structure and inferred population-specific demographic histories, based upon observed levels and patterns of within and between population variation. To provide a comprehensive assessment, we employed several commonly-used model-free (MSMC2 [Schiffels and Durbin 2014; K. Wang et al. 2020] and Stairway Plot2 [Liu and Fu 2020]), and model-based (fastsimcoal2 [Excoffier et al. 2013, 2021; Marchi et al. 2024] and *δaδi* [Gutenkunst et al. 2009]) approaches. To assess the fit of the resulting inference to the empirical data, we used simulation to compare expected versus observed levels and patterns of variation. Finally, we re-estimated model parameters under alternative mutation rates (Spatola et al. 2026) in order to account for underlying uncertainty in these scaling assumptions. Results suggest that the two populations likely diverged approximately ∼140,000 generations ago, and subsequently experienced unique size change histories, reaching current population sizes of ∼220,000 and ∼14,000 individuals for the Chinese and Indian populations, respectively. The correspondingly inferred demographic dynamics fit all assessed aspects of the empirical data, and this inference thus additionally provides a necessary component of a neutral baseline model for future genomic studies ranging from conducting selection inference to quantifying mutational spectra to performing genome-wide association studies (Crisci et al. 2012; Johri et al. 2022a,b; Jensen 2023; Ghafoor et al. 2023; Soni and Jensen 2024; Terbot et al. 2025b; Soni et al. 2025d,2026).

## MATERIALS AND METHODS

### Animal subjects

Rhesus macaques were housed in indoor or outdoor social housing at the Oregon National Primate Research Center (ONPRC). All husbandry practices conducted are performed in accordance with federal guidelines and regulations as stated in the National Institutes of Health Guide for the Care of and Use of Laboratory Animals. ONPRC is accredited by the Association for Assessment and Accreditation of Laboratory Animal Care, International. Buffy coat samples were previously collected and stored under Oregon Health and Science University (OHSU) IACUC protocol #IP00000367.

### Samples and sequencing

We isolated high molecular weight DNA from buffy coat samples of 20 unrelated rhesus macaque (*Macaca mulatta*) individuals from the research colony maintained at the Oregon National Primate Research Center. Of the 20 individuals, ten were of Chinese origin and ten of Indian origin. For each sample, we fragmented the DNA to an approximate size range of 10–20 kb using a Megaruptor 3 (Diagenode, Liège, Belgium), purified the sheared DNA with SMRTbell cleanup beads, and generated sequencing libraries using the SMRTbell Prep Kit 3.0. We then performed size selection on a Pippin HT system (Sage Science, Beverly, MA, USA) with an S1 marker targeting fragments between 10 and 25 kb. We quantified the libraries for each sample using a Qubit HS assay (Invitrogen, Carlsbad, CA, USA) and assessed their fragment size distributions on a Femto Pulse system (Agilent, Santa Clara, CA, USA) before preparing them for sequencing with the PacBio Sequel II Sequencing Kit 3.1 configured for HiFi sequencing. Afterward, we loaded the libraries onto Revio SMRT Cells and sequenced them in CCS mode with 24-hour movie times.

### Variant calling

To avoid artifacts, we converted the raw PacBio HiFi reads to FASTQ format using the *bam2fastq* function implemented in pbtk v.3.4.0 (https://github.com/PacificBiosciences/pbtk) and discarded any reads shorter than 1 kb or containing more than 40% of bases below a Phred-scaled quality score of 20 using fastplong v.0.2.0 (Chen et al. 2018; Chen 2023) with the “*-l* 1000 *-u* 40 *-q* 20” parameters. We then mapped the quality-controlled reads to the rhesus macaque genome assembly (rheMac10; GenBank assembly: GCA_003339765.3; Warren et al. 2020) using minimap2 v.2.26 (Li 2018). We called variants in each individual using the GPU-accelerated version of DeepVariant v.1.6.1 (Poplin et al. 2018) embedded within the NVIDIA Parabricks software suite v.4.4.0-1 (O’Connell et al. 2023), and then combined the individual call sets to jointly genotype variants using GLnexus v.1.4.1 (Yun et al. 2020). We subsequently phased this call set using WhatsHap v.2.3 (Martin et al. 2016).

### Putatively neutral regions

We limited the variant dataset to putatively neutrally evolving regions to circumvent the confounding effects of both direct purifying selection and background selection (see Soni et al. 2025a,b,c). For this purpose, we used the rheMac10 annotations (Warren et al. 2020), consisting of 40,304 protein-coding genes, to mask sites overlapping within 10 kb of exons, following recommendations of Johri and colleagues (2020, 2023) in generating an evolutionary null model. Masking the flanking regions was necessary to account for the effects of selection at linked sites, which have been shown to bias demographic inference (Ewing and Jensen 2014, 2016; Johri et al. 2020, 2021; Charlesworth and Jensen 2021, 2024; and see Johri et al. 2022b). To account for additional regions in the rhesus macaque genome experiencing purifying selection, we excluded any sites located within primate-constrained sequence elements previously identified across 239 species (Kuderna et al. 2024). Finally, we limited our analyses to genomic regions for which sequencing data was available for all 20 individuals included in this study. To this end, we generated an accessibility mask using the *genomecov* function in BEDTools v.2.30.0 (Quinlan and Hall 2010) that excludes any sites covered by fewer than two long-reads per individual.

### Population structure analysis

We assessed population structure within our sampled individuals as the first step in inferring the demographic history of the Indian and Chinese rhesus macaque populations. We examined individual admixture proportions using ADMIXTURE v.1.3.0 (Alexander et al. 2009) across a range of *K* values from 1 to 5, where the *K* value represents the number of ancestral populations. ADMIXTURE analysis was performed on putatively neutral sites, with all 20 autosomes analyzed jointly. To generate input files for ADMIXTURE, we converted the VCF to a binary PED (BED) file using PLINK v.1.9.0 (Purcell et al. 2007; Chang et al. 2015). ADMIXTURE was run with default cross-validation enabled (*--cv*), and the optimal *K* was determined as the value with the lowest cross-validation error (CVE). To further evaluate sample clustering, we performed a principal component analysis (PCA) using PLINK. We subsequently assessed genetic differentiation between Chinese and Indian rhesus macaque populations by estimating genome-wide Weir and Cockerham’s weighted *F_ST_* using VCFtools v.0.1.14 (Danecek et al. 2011).

### Demographic inference

We applied two model-free approaches — MSMC2 (Schiffels and Durbin 2014; K. Wang et al. 2020) and Stairway Plot2 (Liu and Fu 2020) — and two model-based approaches — fastsimcoal2 (Excoffier et al. 2013, 2021; Marchi et al. 2024) and *δaδi* (Gutenkunst et al. 2009) — to infer the population histories of the Indian and Chinese rhesus macaque populations. For the demographic inference methods that require the site frequency spectrum (SFS) as input, we generated folded SFS from the sequence data for each population and a joint SFS for both populations using easySFS v.0.0.1 (https://github.com/isaacovercast/easySFS). We initially assumed a species-specific per-generation mutation rate of 0.58 × 10^−8^ per site (R. Wang et al. 2020; though see the section “Accounting for uncertainty in mutation rates” below), a recombination rate of 0.448 cM/Mb (Xue et al. 2020), and a generation time of 11 years (Xue et al. 2016) for demographic inference.

#### Demographic inference using MSMC2

We first estimated the demographic history of the Chinese and Indian rhesus macaque populations using MSMC2 v.2.1.4. In brief, we used the Perl script *vcf2multihetsep-v0.04.pl* (Terbot et al. 2025a) to generate MSMC2 input files that contained information on the location of putatively neutral variant sites (*-use*) and accessible invariant sites (*-call*) across the genome. Using these input files, and following the population assignment suggested by ADMIXTURE, we inferred the demographic history for each rhesus macaque population in MSMC2 based on all haplotypes (20 per population) and the default time-segment pattern (*-p* 1*2+25*1+1*2+1*3).

#### Demographic inference using Stairway Plot2

We applied Stairway Plot v.2.0b to infer past changes in the effective population sizes (*Nₑ*) of the Indian and Chinese rhesus macaque populations. The Stairway Plot approach assumes an underlying coalescent history for a single panmictic population under an infinite sites model, and allows step-wise population size changes. We randomly selected 67% of all SNPs to construct a training SFS for each replicate (*pct_training* = 0.67) and allowed 200 re-samplings from the SFS (*ninput* = 200). We selected breakpoints following the suggestions in the Stairway Plot2 manual, i.e., at *n*/4, *n*/2, *n\**¾ and *n*-2, with *n* indicating the sample size (*nrand* = 5, 9, 14, 18), and performed analyses separately for each population, including all individuals (10 per population). The median of 200 inferred *Nₑ* curves was taken as the representative demographic history for that population and uncertainty was summarized using the 2.5^th^ and 97.5^th^ percentiles.

#### Demographic inference using fastsimcoal2

To further explore the size-change history, we used fastsimcoal2 v.2.8 with the joint SFS generated from easySFS as input. We defined a total of 13 demographic models for fastsimcoal2 based on information obtained from the population-structure analyses. Models were categorized into four groups: *simple split with no size change* (M1), *two-epoch variable rate migration models with no size change* (M2), *one size-change models with variable rate migration* (M3), and *models with different size-change timing for the Chinese and Indian populations* (M4), with each model group consisting of different migration scenarios (see Supplementary Figure S1 for detailed model information). Across the models, we used the following parameters: the ancestral population size (*N_anc_*), the current Chinese population size (*N_CH_*), the current Indian population size (*N_IN_*), the time of the split of the two populations (*T_DIV_*), the time of the size change for both populations for M3 models (*T_S_*), the time of the size change for the Chinese population in M4 models (*T_CH_*), the time of the size change for the Indian population in M4 models (*T_IN_*), the Chinese population size before the size change at *T_S_* or *T_CH_* (*N_1_*), the Indian population size before the size change at *T_S_* or *T_IN_* (*N_2_*), the time of the migration change for M2 models (*T_M_*), and three different migration matrices for different migration time points (*m1*, *m2*, *m3*). The migration matrices consist of migration rates from the Chinese to the Indian population (*M_Ch2In_*) and from the Indian to the Chinese population (*M_In2Ch_*).

We assigned all parameters log-uniform priors across the models, with effective population sizes (*N_anc_*, *N_CH_*, *N_IN_*, *N_1_*, *N_2_*) ranging from 10^3^ to 10^7^, migration rates in the migration matrices (*m1*, *m2*, *m3*) ranging from 0.01 to 10^-7^, and the split time (*T_DIV_*) ranging from 10^3^ to 10^7^ generations. We kept all other time parameters (*T_S_*, *T_M_*, *T_CH_*, *T_IN_*) between (1-*T_DIV_*) generations. We simulated 250 replicates of each model in fastsimcoal2 with the following parameter settings: *-n* 150,000 (the number of coalescent simulations to perform per replicate), *-L* 100 (the number of expectation maximization cycles), *-M* (performs parameter estimation by maximum composite likelihood from the SFS), *-m* (computes the SFS for the minor allele for each population sample and for the joint SFS for two populations), and *-y* 3 (resets parameters after three cycles without likelihood improvement). We identified the best-supported model and the parameter combinations for each model group based on the maximum likelihood and the minimum difference between the maximum observed likelihood (*MaxObsLhood*) and the maximum estimated likelihood (*MaxEstLhood*).

#### Demographic inference using δaδi

We applied *δaδi* to further validate the demographic estimates for the two rhesus macaque populations. We first performed optimizations using the dadi_pipeline v.3.1.6 (Portick 2017). For the initial optimization, we included 20 demographic models in the *Two-Population Pipeline* (https://github.com/dportik/dadi_pipeline/tree/master/Two_Population_Pipeline), which include simple models, simple models plus instantaneous size changes, ancient migration, secondary contact, ancient migration or secondary contact plus instantaneous size change, two-epoch models with continuous migration, and three-epoch models with migration and instantaneous size change variations. We inferred the best-fit model as the model with the consistently highest likelihood and lowest Akaike information criterion (AIC) score (Akaike 1974) across multiple runs.

We then re-estimated the parameters by running 100 *δaδi* simulations on the inferred best model (based on the lowest AIC values across multiple runs), *sym_mig_size* (from the dadi_pipeline; Portick 2017), which comprises a divergence of two populations with continuous symmetrical migration followed by an instantaneous size change in both populations. The model consists of seven parameters: *nu1a* (the ratio of the population size of the Chinese population relative to the ancestral population after the population split), *nu2a* (the ratio of the population size of the Indian population relative to the ancestral population after the population split), *nu1b* (the ratio of the population size of the Chinese population relative to the ancestral population after instantaneous size change), *nu2b* (the ratio of the population size of the Indian population relative to the ancestral population after instantaneous size change), *T1* (the time of population split in units of *2*N_a_* generations, where *N_a_* is the effective size of the ancestral population), *T2* (the time of instantaneous population size change in units of *2*N_a_* generations), and *M* (the migration rate between populations measured as *2*N_a_*m*). In these simulations, we set the optimization function to 300 maximum iterations, using starting parameter values of *nu1a*=1, *nu2a*=1, *nu1b*=1, *nu2b*=1, *m*=0.01, *T1*=0.5, and *T2*=0.5. We set the parameters to adjust two-fold from the initial set value using *δaδi’*s *perturb_params* function with the upper and lower bounds set in the ranges as follows: 0.01 ≤ *nu1a* ≤ 100, 0.01 ≤ *nu2a* ≤ 100, 0.01 ≤ *nu1b* ≤ 100, 0.01 ≤ *nu2b* ≤ 100, 0.001 ≤ *m* ≤ 0.1, 0.01 ≤ *T1* ≤ 10, 0.001 ≤ *T2* ≤ 10. We selected the simulation replicate with the highest log-likelihood as the best-fit parameter combination, and calculated the corresponding demographic values based on these best-fit parameters.

### Demographic model validation using msprime simulations

We assessed the fit of the best-fit models and their parameters from each demographic approach to the empirical data via simulations using msprime v.1.3.2 (Baumdicker et al. 2022). Here, we evaluated seven demographic models: one best-fitting model from each of MSMC2, Stairway Plot2, and *δaδi*, and four from fastsimcoal2, each representing the best-fit model from one of the four predefined fastsimcoal2 model groups (M1-M4). We carried out the msprime simulations per autosome, with each autosome simulated 10 times (for a total of 200 simulations per model). The recombination rate was set to 0.448 cM/Mb (Xue et al. 2020), and the mutation rate was set to 0.58 × 10^−8^ per site per generation (R. Wang et al. 2020) to match previously used values. For each simulated model, we generated the folded SFS using easySFS and calculated per chromosome pairwise Weir and Cockerham’s weighted *F_ST_* values using VCFtools v.0.1.14, matching the summary statistics computed from the empirical data. We then compared the estimated SFS and mean *F_ST_* values — averaged across per-chromosome values — for each msprime-simulated model with the observed empirical values to assess the fit of each demographic model to the data. We considered the demographic model indicating the closest agreement across summary statistics to be the best-supported demographic scenario.

Furthermore, we assessed the fit of empirically-observed genome-wide linkage disequilibrium — a data summary not utilized for these model-fitting procedures — to that expected under the best-fitting model. To this end, we estimated pairwise *r^2^* values (i.e., the squared correlation of pairwise allele frequencies) separately for each population using PLINK v.1.9.0 within a 1 Mb sliding window (*--ld-window-kb* 1000). We grouped pairwise *r^2^* values into distance bins based on the physical distance between SNP pairs, and calculated the mean *r^2^* for each bin. We then calculated the chromosome-level linkage disequilibrium estimates by averaging the mean *r^2^* values across all bins. For simulated datasets, we applied the same procedure independently to each of the 10 simulation replicates for both populations, and calculated the final chromosome-level estimates by averaging across replicates.

### Accounting for uncertainty in mutation rates

To investigate the robustness of the best-supported demographic model, we also evaluated the fit for a model with alternative mutation rate estimates (see Soni et al. 2024b). Specifically, we simulated 250 replicates of the best-fit model (*M4_consMig*) in fastsimcoal2, keeping the same parameter options as in the initial fastsimcoal2 runs (see the section “Demographic inference using fastsimcoal2”) but re-estimating under a species-specific per-generation mutation rate of 1.49 × 10^−8^ per site (Spatola et al. 2026). We then simulated parameter values from the maximum likelihood replicate using msprime. We performed the msprime simulations as described above with 10 simulations per autosome (see the section “Demographic model validation using msprime simulations”). We generated the simulated SFS using easySFS and re-estimated the mean *F_ST_* from the simulated SFS using VCFtools. We then compared the parameter estimates from this msprime simulation with both those obtained under the previous per-generation mutation rate of 0.58 × 10^−8^ per site and those obtained from the empirical data.

## RESULTS

### Population genomic data

We generated whole genome PacBio HiFi sequencing data for 20 rhesus macaque individuals — ten of Chinese origin and ten of Indian origin — and mapped the resulting long-reads against the rhesus macaque reference assembly (rheMac10; GenBank assembly: GCA_003339765.3; Warren et al. 2020). From these mappings, we identified variants using DeepVariant (Poplin et al. 2018), a deep neural network approach previously demonstrated to exhibit high precision-recall rates in humans (>0.99; Kolesnikov et al. 2024). Using this approach, we identified 36 million autosomal, biallelic SNPs (Ts/Tv = 2.2) that were genotyped in all individuals. We identified regions on the autosomes (chromosomes 1–20) for which sequencing data was available for all 20 individuals included in this study; this accessibility mask excluded 49.3% of the rhesus macaque genome. Additionally, in order to circumvent the confounding effects of both direct purifying selection and background selection, we limited this variant dataset to putatively neutrally evolving regions, excluding any sites within 10 kb of exons (Warren et al. 2020) or sequence elements constrained across the primate clade (Kuderna et al. 2024). Demographic analyses focused on the 8.5 million autosomal, biallelic SNPs (Ts/Tv = 2.1) located in accessible, putatively neutral regions.

### Population structure analysis

As the initial step in demographic inference, we estimated the number of populations implied by the empirical data using ADMIXTURE. The lowest CVE values corresponded to both one- and two-population assignments in that they were nearly identical (*K*=1, CVE=0.63371; *K*=2, CVE=0.63491; *K*=3, CVE=0.79136; *K*=4, CVE=1.00824; *K*=5, CVE=1.11946). We visually evaluated the individual admixture proportion bar plots, which revealed a consistent subdivision pattern, with each individual assigned 100% to one of the two populations at *K* = 2 (Figure 1; and see Supplementary Figure S2 for *K* = 1–5). A PCA further confirmed the population assignment to two clusters, with a clear separation in PC1, which explained 13.1% of the variance (Supplementary Figure S3). After evaluating the ADMIXTURE and PCA results, and observing an average *F_ST_* value between the Chinese and Indian rhesus macaque populations across autosomes of 0.1476, we considered *K*=2 to represent the most optimal assignment for demographic history inference.

**Figure 1:**
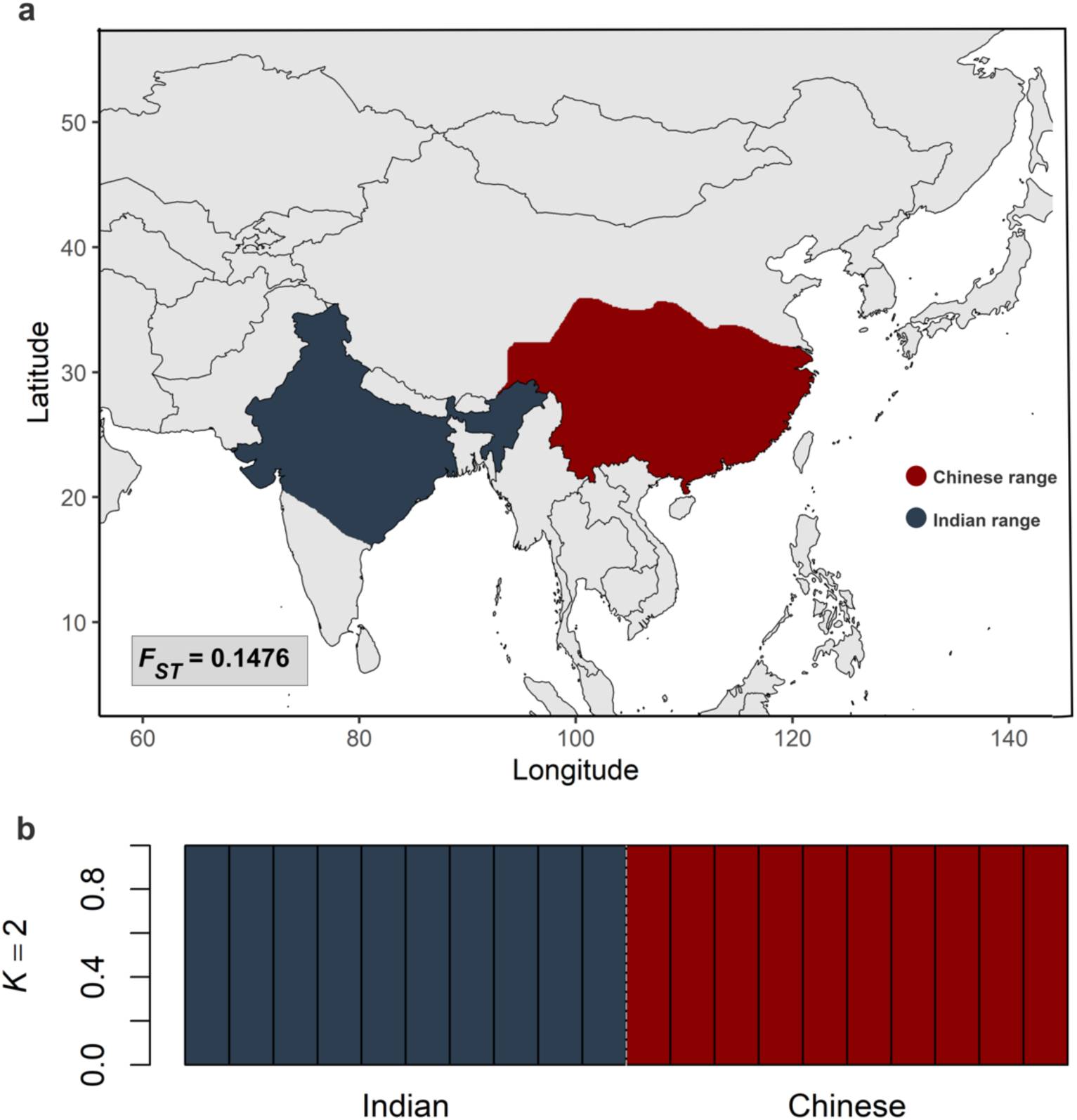
Geographic distribution and genetic structure of rhesus macaque (*Macaca mulatta*) populations in China and India. (a) Natural geographic range of rhesus macaque populations in China (shown in red) and India (blue) (obtained from the IUCN 2025). The empirically estimated *F_ST_* value displayed in the bottom-left corner indicate the level of genetic differentiation observed between the Chinese and Indian populations used in this study. (b) Population structure inferred using ADMIXTURE at *K* = 2. Each vertical bar represents one individual, and colors denote inferred ancestry proportions (with the Chinese population shown in red and the Indian population shown in blue). ADMIXTURE results for *K* = 1–5 are provided in Supplementary Figure S2.

### Demographic inference

#### Demographic inference using MSMC2

The demographic history estimated using MSMC2 was characterized by population decline and subsequent recovery in both rhesus macaque populations (Figure 2a). The ancestral effective size of the Chinese population was estimated at ∼120,000 individuals, and population decline was observed until approximately 10,000 years ago. Subsequently, the Chinese population began to recover, but a rapid decline was detected about 800 years ago, followed by growth in recent years. The estimated current effective size of the Chinese population was ∼30,000 individuals. The ancestral effective size of the Indian population was estimated at ∼100,000 individuals and followed a similar pattern to the Chinese population until roughly 10,000 years ago. The effective size of the Indian population during the recovery phase exceeded that of the Chinese, reaching a peak of ∼70,000 individuals. The recent decline in population size suggests that the current effective size in Indian rhesus macaques is ∼40,000 individuals. The visual examination of the *N_e_* trajectories for each population implies population divergence around 10,000 years ago.

**Figure 2:**
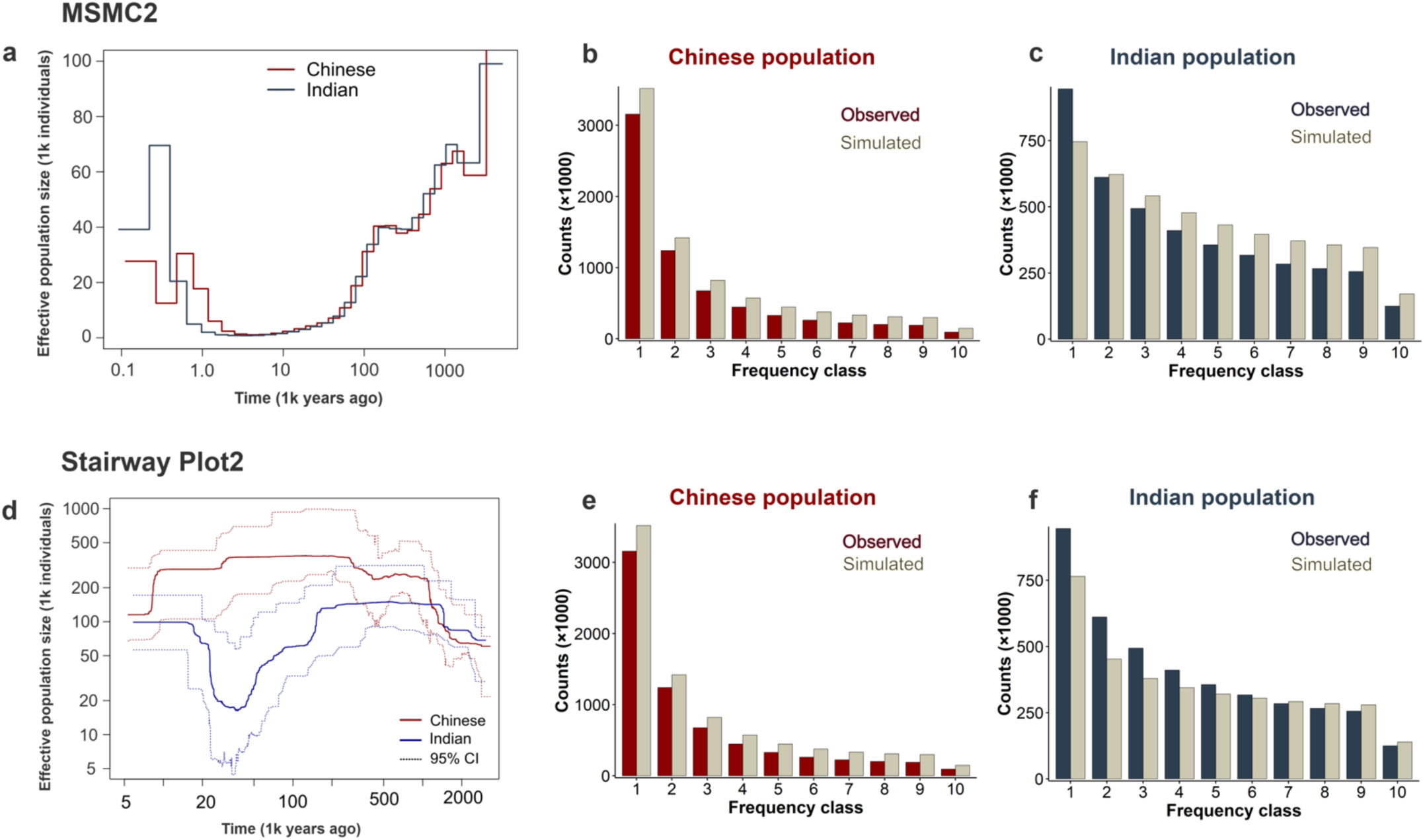
Demographic inference using MSMC2 and Stairway Plot2. (top) Demographic inference using MSMC2. (a) Diagram of the best-fitting MSMC2 demographic model. (b-c) Folded SFS for the Chinese (red) and Indian (blue) populations compared to the best-fitting MSMC2 model (shown in gray). (bottom) Demographic inference using Stairway Plot2. (d) Diagram of the best-fitting Stairway Plot2 demographic model. (e-f) Folded SFS for the Chinese (red) and Indian (blue) populations compared to the best-fitting Stairway Plot2 model SFS (shown in gray). Note the differing scaling on the x- and y-axes reported by the two approaches.

We evaluated the fit of the model estimated by MSMC2 to the data using msprime simulations and found that the model fit to the observed data was poor, as shown by the simulated SFS compared with the empirically-observed SFS (Figures 2b and 2c, respectively). In the Chinese population, the simulated SFS underestimated counts of singletons and doubletons and overestimated counts of intermediate and high-frequency alleles (Figure 2b). A similar pattern was observed in the Indian population where singletons were underestimated while all other frequency classes were overestimated (Figure 2c). Additionally, *F_ST_* estimated from the simulated SFS (0.0836) was substantially lower than that observed in the empirical data (0.1476), suggesting an underestimation in population differentiation.

#### Demographic inference using Stairway Plot2

Stairway Plot2, a composite-likelihood method that infers changes in *N_e_* over time from empirically-observed SFS, suggested distinct demographic histories for the Chinese and Indian rhesus macaque populations. In the Chinese population, the inferred *Nₑ* trajectory indicated a period of population growth beginning approximately 2 million years ago, reaching a maximum *Nₑ* of ∼300,000 individuals around 250,000 years ago (Figure 2d). This was followed by a long period of relative demographic stability with little change in population size. A subsequent decline began roughly 10,000 years ago, leading to a recent *Nₑ* of ∼110,000 individuals. The stairway plot for the Indian population suggests that population growth began around 2 million years ago, reaching a peak *Nₑ* of ∼130,000 individuals about 1.5 million years ago. The population remained stable for an extended period before experiencing a marked demographic contraction starting around 150,000 years ago, during which *Nₑ* declined to ∼20,000 individuals. The Indian population recovery started roughly 30,000 years ago and peaked about 15,000 years ago. The population maintained a relatively consistent size up to the present, with a contemporary *Nₑ* of ∼100,000 individuals.

We used msprime simulations to assess the fit of the model estimated by Stairway Plot2 to the data. The simulated SFS for the Chinese population overestimated allele counts across all frequency bins with a larger deviation for rare alleles than for more frequent alleles (Figure 2e). In contrast, the simulated SFS of the Indian population more closely matched the observed distribution across most frequency classes particularly at high frequencies (Figure 2f). However, *F_ST_* estimated from the simulated SFS (0.2396) was substantially higher than that observed in the empirical data (0.1476), indicating an overestimation of differentiation between Chinese and Indian rhesus macaque populations.

#### Demographic inference using fastsimcoal2

We explored a set of 13 demographic models categorized into four general model groups — *simple split models with no size change* (M1), *two-epoch migration models with no size change* (M2), *one size-change models with migration* (M3), and *models with different size-change timing for the Chinese and Indian populations* (M4) — using fastsimcoal2 (Supplementary Figure S1; Supplementary Table S1). Within each group, we identified the best-fitting model and corresponding parameters using the minimum difference between the maximum observed and estimated likelihoods (*MaxObsLhood* – *MaxEstLhood*) across 250 independent replicates (Supplementary Table S2).

The best *simple split with no size change* model in group M1 (*M1_Mig*) supported a split between the Chinese and Indian populations around 118,943 generations ago, with continuous asymmetric migration between the populations. Migration from the Indian population to the Chinese population (*M_In2Ch_* = 2.53e-05) was estimated to be higher than that from the Chinese population to the Indian population (*M_Ch2In_* = 1.14e-06). Population size estimates indicated an ancestral size (*N_anc_*) of 64,255 individuals. In the post-split period, population sizes remained unchanged, with current sizes of the Chinese population (*N_CH_*) at 275,490 individuals and the Indian population (*N_IN_*) at 22,870 individuals.

The best-fitting *two-epoch migration with no size change* model in group M2 (*M2_difMig*) indicated an initial divergence of the Chinese and Indian rhesus macaque populations around 133,874 generations ago from an *N_anc_* of 60,007 individuals. Following the split, the two populations exchanged migrants at asymmetric rates (*M_Ch2In_* = 2.53e-06; *M_In2Ch_* = 2.11e-06). Approximately 5,878 generations ago, the patterns of migration shifted to a second epoch, again characterized by asymmetric migration rates between the populations (*M_Ch2In_* = 4.07e-07; *M_In2Ch_* = 3.68e-05). Since the population split, throughout the migration phases, the population sizes remained unchanged, estimated as 265,069 individuals in the Chinese population (*N_CH_*) and 27,010 individuals in the Indian population (*N_IN_*).

The best-supported *one size-change with migration* model in group M3 (*M3_consMig*) implied continuous migration from the population split to the present, with asymmetric gene flow between the populations (*M_Ch2In_* = 3.49e-06; *M_In2Ch_* = 1.69e-05). This model supported the divergence of the Chinese and Indian populations approximately 104,615 generations ago from an *N_anc_* of 68,473 individuals, resulting in population sizes of 259,595 and 102,844 individuals for the Chinese (*N_CH_*) and Indian (*N_IN_*) populations, respectively. Around 7,053 generations ago, both populations underwent a size change, resulting in the current population sizes of 268,928 individuals in the Chinese population (*N_CH_*) and 14,203 individuals in the Indian population (*N_IN_*).

The highest-likelihood *different time size change* model in group M4 (*M4_consMig*) suggested divergence 139,371 generations ago from an *N_anc_* of 64,723 individuals (Figure 3a). At the time of the split, the Indian population expanded to a population size of 270,072 individuals, while the Chinese population remained at a population size of 76,025 individuals. Approximately 71,781 generations ago, the substantial expansion of the Chinese population enabled it to reach its current size of 219,783 individuals (*N_CH_*). The Indian population underwent a notable decline in size 8,814 generations ago, reaching the current size of 14,091 individuals (*N_IN_*). The ongoing asymmetric gene flow since the split of the two populations demonstrated a higher migration rate from the Indian to the Chinese population than the reverse (*M_Ch2In_* = 6.63e-06; *M_In2Ch_* = 1.46e-05), consistent with the estimated migration matrices from the best-fitting models in other groups.

**Figure 3:**
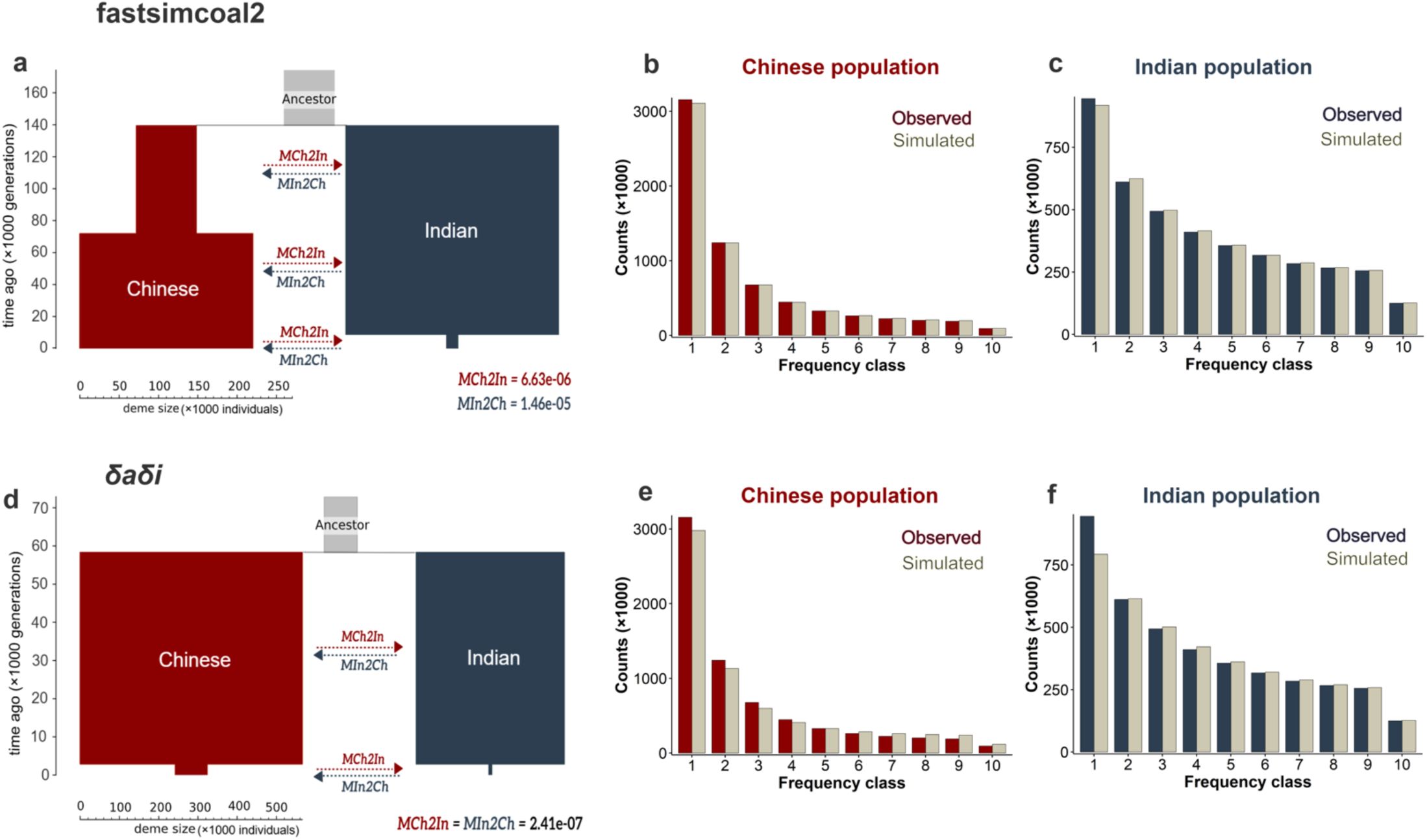
Demographic inference using fastsimcoal2 and *δaδi*. (top) Demographic inference using fastsimcoal2. (a) Diagram of the best-fitting fastsimcoal2 demographic model. (b-c) Folded SFS for the Chinese (red) and Indian (blue) populations compared to the best-fitting fastsimcoal2 model (shown in gray). (bottom) Demographic inference using *δaδi*. (d) Diagram of the best-fitting *δaδi* demographic model. (e-f) Folded SFS for the Chinese (red) and Indian (blue) populations compared to the best-fitting *δaδi* model (shown in gray). Note the differing scaling on the axes between the two approaches.

The highest-likelihood model across all four groups was observed for the *M4_consMig* model (*MaxObsLhood – MaxEstLhood* = 2959.77; Supplementary Table S2); nevertheless, we evaluated the fit of the best-supported model from each of the four model groups (M1-M4). For the Chinese population, the best-fit models from all groups except M2 produced simulated SFS that closely matched the empirically-observed SFS (see Figure 3b for the highest-likelihood model, *M4_consMig*, and Supplementary Figures S4a-d for the best-supported model from each of the four model groups). These models yielded broadly consistent estimates of the current population sizes (*N_CH_* and *N_IN_*), ancestral population size (*N_anc_*), and split time (*T_DIV_*) (Supplementary Table S1). In contrast, the best-fit model from group M2 (*M2_difMig*) overestimated the number of singletons. For the Indian population, the simulated SFS differed across the four groups and the estimated model of group M4 (*M4_consMig*) produced the best-fitting SFS (see Figure 3c for the highest-likelihood model, *M4_consMig*, and Supplementary Figures S4 e-h for the best-supported model from each of the four model groups). Moreover, *F_ST_* estimated from the *M4_consMig* model (0.1498, Supplementary Table S3) was consistent with that observed from the empirical data (0.1476).

As this best-fitting model was found to match both empirically-observed SFS and between-population divergence, we further assessed the fit of this model to empirically-observed, genome-wide linkage disequilibrium patterns. This comparison is of particular value given that summaries of linkage disequilibrium were not used in the fastsimcoal2 model-fitting procedure and thus this comparison represents a semi-independent assessment of fit. As shown in Table 1, the best-fitting demographic model as inferred via the SFS also well-predicts empirically-observed values of linkage disequilibrium.

**Table 1:** Chromosome-level, per-population linkage disequilibrium (*r^2^*) values compared between empirical data and simulated data based on the best-fitting demographic model.

| chromosome | China |  | India |  |
| --- | --- | --- | --- | --- |
|  | empirical | simulations | empirical | simulations |
| 1 | 0.11275 | 0.11198 | 0.12817 | 0.12066 |
| 2 | 0.11248 | 0.11199 | 0.12525 | 0.12120 |
| 3 | 0.11152 | 0.11227 | 0.12810 | 0.12155 |
| 4 | 0.11310 | 0.11207 | 0.12653 | 0.12171 |
| 5 | 0.11219 | 0.11195 | 0.12673 | 0.12108 |
| 6 | 0.11361 | 0.11195 | 0.13040 | 0.12156 |
| 7 | 0.11198 | 0.11197 | 0.12532 | 0.12111 |
| 8 | 0.11164 | 0.11236 | 0.12476 | 0.12136 |
| 9 | 0.11321 | 0.11197 | 0.12350 | 0.12101 |
| 10 | 0.11293 | 0.11173 | 0.12331 | 0.12219 |
| 11 | 0.11276 | 0.11235 | 0.12538 | 0.12084 |
| 12 | 0.11307 | 0.11197 | 0.12477 | 0.12121 |
| 13 | 0.11320 | 0.11226 | 0.13071 | 0.12183 |
| 14 | 0.11257 | 0.11220 | 0.12557 | 0.12193 |
| 15 | 0.11319 | 0.11190 | 0.12671 | 0.12151 |
| 16 | 0.11276 | 0.11186 | 0.12003 | 0.12090 |
| 17 | 0.11244 | 0.11218 | 0.12298 | 0.12135 |
| 18 | 0.11252 | 0.11245 | 0.12870 | 0.12067 |
| 19 | 0.11146 | 0.11171 | 0.12172 | 0.11917 |
| 20 | 0.11252 | 0.11212 | 0.12098 | 0.12095 |

#### Demographic inference using δaδi

We performed initial optimizations using the dadi_pipeline across 20 demographic models (see “Materials and Methods” for details). The *sym_mig_size* was inferred as the best-fit model with the highest log-likelihood (-45,427.15) and the lowest AIC (90,868.3) scores across five rounds of optimization. To infer the best-fit parameter values from the *sym_mig_size* model, we performed 100 independent *δaδi* simulations (Supplementary Figure S5). Under this model, an ancestral population of size 84,133 was split into two populations approximately 58,330 generations ago, giving rise to Chinese and Indian lineages with sizes of 565,749 and 377,188 individuals, respectively (Figure 3d). Both populations experienced an instantaneous decline in size 2,894 generations ago, resulting in the current population sizes of 80,960 and 7,247 individuals for the Chinese and Indian population, respectively. Migration continued throughout the post-split period with a symmetrical rate of 2.41e-07.

We evaluated the fit of the best model inferred using *δaδi*, *sym_mig_size* to the data using msprime simulations. The simulated SFS for the Chinese population underestimated low-frequency alleles while overestimating high-frequency alleles (Figure 3e). For the Indian population, the simulated SFS was much closer to the observed SFS across all classes except for singletons, where it was underestimated (Figure 3f). Despite the subtle mismatches in the SFS, the estimated *F_ST_* (0.1435) from the simulated SFS was consistent with the empirically-observed *F_ST_* (0.1476).

### Accounting for uncertainty in the mutation rate

To account for uncertainty in the mutation rate estimates used for scaling, we evaluated the best-fit model under the alternate scaling of 1.49 × 10^−8^ per site per generation using 250 independent replications performed in fastsimcoal2 (Spatola et al. 2026). The simulation with the highest likelihood (*MaxObsLhood – MaxEstLhood* = 1,785.89) suggested a population split around 80,351 generations ago from an ancestral population of 21,986 individuals (Supplementary Figure S6a). At the time of the split, the size of the Chinese population was 24,710 individuals while the Indian population expanded to 45,917 individuals. With this alternative mutation rate scaling, the expansion in the Chinese population occurred 31,538 generations ago compared to 71,781 generations ago under the previously used mutation rate (0.58 × 10^−8^ per site per generation), reaching a current size of 97,535 individuals (Supplementary Figure S6b). A substantial decline in the Indian population was observed 2,884 generations ago, reaching a current size of 5,601 individuals, compared to 8,814 generations ago under the alternative mutation rate scaling (Supplementary Figure S6c). The ongoing asymmetric gene flow since the split of the two populations was fit by a higher migration rate from the Indian to the Chinese population (*M_Ch2In_* = 1.16e-05; *M_In2Ch_* = 4.10e-05), consistent with the previously estimated migration matrix from the best-fit model.

We evaluated the fit of the parameter values from the best model inferred by fastsimcoal2 under this alternative mutation rate with msprime. As expected, given that the alternative mutation rate simply represents a change in scaling, the simulated SFS continues to closely match the observed distribution across all frequency classes in both populations (Supplementary Figure S6b-c), and the level of population differentiation remains consistent as well (with an *F_ST_* estimated from the simulated SFS of 0.1493).

## DISCUSSION

To infer the demographic histories of the Indian and Chinese rhesus macaque populations, we first assessed population structure using ADMIXTURE, followed by two non-model-based approaches (MSMC2 and Stairway Plot2) and two model-based approaches (fastsimcoal2 and *δaδi*). For each demographic inference method, we assessed the fit of the best-supported model by performing simulations under the inferred demographic parameters and compared the simulated SFS and *F_ST_* values with those observed in the empirical data. The fit of the SFS resulting from both the estimated MSMC2 and Stairway Plot2 models to the observed data was poor, and those models can thus be discarded. Both fastsimcoal2 and *δaδi* produced well-fitting models, with the fastsimcoal2 model*, M4_consMig*, identified as the best-supported model in recapitulating the empirically-observed SFS and *F_ST_* values. Inference under this model suggested that the Chinese and Indian populations diverged from an ancestral population of 64,723 individuals roughly 140,000 generations ago. Both populations underwent significant size changes at different times, with notable changes around 72,000 generations ago for the Chinese population and around 9,000 generations ago for the Indian population. The current sizes were estimated at 219,783 individuals in the Chinese population and 14,091 individuals in the Indian population, consistent with the greater levels of genetic variation observed in the former. Throughout the post-split period, a consistent yet asymmetrical migration rate was inferred (*M_Ch2In_* = 6.63e-06; *M_In2Ch_* = 1.46e-05).

Given the strongly differing performance amongst methods (Supplementary Table S4), it is important to consider the various underlying details of the methodologies. MSMC2 is based on the Sequentially Markovian Coalescent (SMC) framework (Marjoram and Wall 2006), which models recombination to infer changes in the coalescence rate over time in order to estimate the time to the most recent common ancestor between pairs of haplotypes across multiple genomes (Schiffels and Durbin 2014; and see Beichman et al. 2018). Stairway Plot2 reconstructs historical changes in effective population size using a composite likelihood based on the given SFS under a coalescent framework (Liu and Fu 2020). Since both MSMC2 and Stairway Plot2 assume panmixia and do not explicitly model migration, we applied these methods separately to each population. Notably, the poor resulting fit of these models to the empirical data is consistent with previous findings; for example, MSMC2 has been shown to overestimate *N_e_* in recent time intervals (Mazet et al. 2016; Chikhi et al. 2017; Beichman et al. 2017; Hilgers et al. 2025), suggest false size changes in the presence of population structure (Orozco-terWengel 2016), incorrectly infer growth prior to instantaneous bottlenecks (Bunnefeld et al. 2015), and generally perform poorly even under constant population sizes in addition to being highly sensitive to the amount of data utilized for analysis (Johri et al. 2021). Together, these factors are likely explanatory of the mismatch between the model estimates and the empirical data for these two nonparametric approaches. Moreover, this accumulated body of work strongly suggests that these approaches are much more widely used than is justified by their performance, and once again highlights the great importance of taking the additional steps of evaluating the fit of estimated models to the observed data being analyzed when performing such inference (Johri et al. 2022a).

The two model-based approaches employed both rely on fitting the observed SFS to evaluate predefined demographic scenarios. fastsimcoal2 utilizes a composite-likelihood framework to compare alternative demographic models by simulating expected SFS across different parameter combinations and identifying the best-fitting model (Excoffier et al. 2013, 2021; Marchi et al. 2024). *δaδi* uses a diffusion approximation to model the expected SFS under predefined demographic models and then compares this expected SFS with the observed data to infer underlying demographic parameters (Gutenkunst et al. 2009). Compared with model-free approaches, the estimated models from fastsimcoal2 and *δaδi* provided greatly improved fits to both the empirically-observed SFS and *F_ST_* values. However, the demographic histories inferred by fastsimcoal2 and *δaδi* were fairly distinct, as has been described in previous studies (e.g., Laurent et al. 2016; Terbot et al. 2026). Although both methods rely on the SFS and explicit modeling, they differ in likelihood approximation, parameter scaling, and optimization, which will contribute to these downstream differences. However, it is naturally the case generally when performing statistical inference that multiple different models may all provide satisfactory fits to the data; for this reason, population genetic analysis of this sort is best viewed as a narrowing down of viable hypotheses rather than as a means of identifying a single ‘true model’ (Johri et al. 2022a). Though both reasonably well-fitting, in this application notable differences were observed between the fit of the resulting fastsimcoal2 and *δaδi* models (Figure 3) that allowed for their differentiation, and for the quantification of the best-supported model. Moreover, given that this high-quality long-read sequencing data additionally provides for an accurate empirical measure of genome-wide patterns of linkage disequilibrium, this allows for an additional means of model assessment that is independent of the inference procedure itself. As such, the fit between empirically-observed and model-predicted patterns of linkage disequilibrium provides further support for the *M4_consMig* model (Table 1).

Assuming a generation time of 11 years (Xue et al. 2016), the population split in our model occurred approximately 140,000 generations ago, dating to ∼1.54 million years ago. This timing roughly corresponds to the Yuanmu uplift ∼1.6 million years ago during the Early Pleistocene, which contributed to the creation of greater elevations in the Himalaya (Zheng et al. 2002). This uplift likely intensified regional cooling, glaciation and drainage reorganization, potentially altering habitats and reinforcing geographic barriers (including new river systems) separating the Chinese and Indian populations. Moreover, the large expansion in the Chinese population, inferred ∼790 thousand years ago, coincides with the end of the Xixiabangma Glaciation (∼1,200–800 thousand years ago), one of the major Early Pleistocene glacial periods on the Tibetan Plateau. During this time, climate changes and an intensified monsoon likely expanded suitable habitats and increased resource availability, promoting the expansion of rhesus macaques in this region.

Rhesus macaque population size changes in China during glacial and post-glacial periods have been previously reported (Liu et al. 2018; Zhou et al. 2024; Terbot et al. 2025b). Finally, the severe decline in the Indian population ∼97 thousand years ago coincides with the weakening of the Indian Summer Monsoon during interglacial substages MIS 5c–5a (∼104-82 thousand years ago; Band et al. 2022), which is believed to have reduced forest coverage and habitat connectivity across the Indian subcontinent. Taken together, the glaciation and tectonic uplift that led to regional environmental changes likely contributed to the split and subsequent contrasting demographic histories of these populations.

Previous studies have also inferred the divergence time of Indian and Chinese rhesus macaques using different approaches, including nuclear and mitochondrial DNA analyses (Smith and McDonough 2005; Hernandez et al. 2007; Hasan et al. 2014; Xue et al. 2016; Zhou et al. 2024), as well as evidence from climate change, sea-level fluctuations, and paleontological records (Abegg and Thierry 2002). Most notably, a previous population genetic analyses of 1,467 SNPs across five Encyclopedia of DNA Elements (ENCODE) regions on separate autosomes suggested a threefold expansion and a fourfold contraction in the Chinese and Indian populations, respectively (Hernandez et al. 2007). Their inferred ancestral and current population sizes are consistent with our estimates (*N_anc_* = 64,723 vs. ∼73,070; *N_CH_* = 219,783 vs. ∼239,704; *N_In_* = 14,091 vs. ∼17,014). However, our estimated divergence time of ∼140,000 generations ago is much older than their suggested divergence of ∼162,000 years ago. Importantly, our whole-genome, high-fidelity, long-read data analyzed here (consisting of 8.5 million autosomal SNPs), allowed for the exploration of more complex population models including, for example, a variety of gene flow scenarios. As we found strong statistical support for limited but long-term gene flow, it is to be expected that the incorporation of these parameters will result in older split times relative to models that do not incorporate the possibility of gene flow. Furthermore, as shown, our resulting estimates well-predict not only population-specific SFS, but also the observed level of population differentiation and linkage disequilibrium.

Taken together, this analysis represents the most complete characterization of the demographic history of Chinese and Indian rhesus macaque populations to date, utilizing the highest-quality data resource for the species available. These histories will provide critical baseline models for future population genomic analyses ranging from the estimation of fine-scale rates and patterns of population-specific recombination to performing scans for genes recently targeted by the action of positive or balancing selection. Importantly, these results also demonstrate the notable differentiation between these two populations, and the differing population histories characterizing them, which in turn has resulted in differing underlying levels of population variation and haplotype structure; considerations that also must necessarily be accounted for in a population-specific manner when performing association studies in existing biomedical populations containing variable Chinese and Indian ancestry.

## ACKNOWLEDGEMENTS

We would like to thank Sam Peterson and the team at the Oregon National Primate Research Center (ONPRC) for providing the rhesus macaque samples used in this study. DNA extraction was performed at the ONPRC (Beaverton, OR, USA), library preparation and PacBio HiFi sequencing were performed at the Arizona Genomics Institute at the University of Arizona (Tucson, AZ, USA). Computations were performed on the Sol supercomputer at Arizona State University (Jennewein et al. 2023).

## FUNDING

This work was supported by the National Institute of General Medical Sciences of the National Institutes of Health under Award Number R35GM151008 to SPP and Award Number R35GM139383 to JDJ, as well as the ONPRC NIH base grant P51OD011092 and the ONPRC Primate Genetics Core (RRID:SCR_027583). CJV was supported by the National Science Foundation CAREER Award DEB-2045343 to SPP. The content is solely the responsibility of the authors and does not necessarily represent the official views of the funders.

## Notes

### Competing Interest Statement

The authors have declared no competing interest.

