## Supplementary Materials for "Inferring the demographic history of Chinese and Indian rhesus macaque (*Macaca mulatta*) populations from PacBio HiFi long-read sequencing data"

### Supplementary Information: Tables and Figures

**Table S1:** The estimated best parameter values for each of 13 demographic models tested with fastsimcoal2, categorized into four groups: *simple split with no size change* (M1), *two-epoch migration models with no size change* (M2), *one size-change models with migration* (M3), and *models with different size-change timing for the Chinese and Indian populations* (M4) (and see Supplementary Figure S1 for detailed model information). Estimated parameters include the ancestral population size ($N_{anc}$), the current Chinese population size ($N_{CH}$), the current Indian population size ($N_{IN}$), the time of the split of two populations ($T_{DIV}$), the time of the size change for both populations for M3 models ($T_S$), the time of the size change for the Chinese population in M4 models ($T_{CH}$), the time of the size change for the Indian population in M4 models ($T_{IN}$), the Chinese population size before the size change at $T_S$ or $T_{CH}$ ($N_1$), the Indian population size before the size change at $T_S$ or $T_{IN}$ ($N_2$), and the time of migration change for M2 models ($T_M$). Migration rates are defined as $M_{Ch2In}$ (from the Chinese to the Indian population) and $M_{In2Ch}$ (from the Indian to the Chinese population). These rates are presented according to the number of migration matrices included: $M_{Ch2In}/M_{In2Ch}$ (for a single migration matrix, *m1*), $M_{Ch2In}/M_{In2Ch}$ and $M_{Ch2In1}/M_{In2Ch1}$ (for two migration matrices, *m1* and *m2*), and $M_{Ch2In}/M_{In2Ch}$, $M_{Ch2In1}/M_{In2Ch1}$ and $M_{Ch2In2}/M_{In2Ch2}$ (for two migration matrices, *m1* and *m2*). Parameters not included in a given model are indicated by a "–" symbol. The parameters obtained from the best model, *M4_consMig*, are highlighted in green. Additionally, the best model was rerun using an alternative mutation rate of 1.49 × $10^{-8}$ per site per generation to account for uncertainty in the mutation rate; this model is referred to as *M4_consMig**.

| **model group** | **model** | **population size** | | | | | **time of split** | **time of population size change** | **migration rate** |
|---|---|---|---|---|---|---|---|---|---|
| | | $N_{CH}$ | $N_{IN}$ | $N_{anc}$ | $N_1$ | $N_2$ | $T_{DIV}$ | $T_M$, $T_S$, $T_{CH}$, or $T_{IN}$ | $M_{Ch2In}$, $M_{In2Ch}$, $M_{Ch2In1}$, $M_{In2Ch1}$, $M_{Ch2In2}$, or $M_{In2Ch2}$ |
| M1 | *M1_noMig* | 416,215 | 55,957 | 105,202 | – | – | 35,342 | – | – |
| | *M1_Mig* | 275,490 | 22,870 | 64,255 | – | – | 118,943 | – | 1.14e-06 ($M_{Ch2In}$)<br>2.53e-05 ($M_{In2Ch}$) |
| M2 | *M2_ancMig* | 273,931 | 24,997 | 65,352 | – | – | 117,623 | 4,260 ($T_M$) | 1.26e-06 ($M_{Ch2In}$)<br>3.74e-05 ($M_{In2Ch}$) |
| | *M2_recMig* | 261,341 | 28,325 | 66,524 | – | – | 117,398 | 98,669 ($T_M$) | 1.85e-06 ($M_{Ch2In}$)<br>2.00e-05 ($M_{In2Ch}$) |
| | *M2_difMig* | 265,069 | 27,010 | 60,007 | – | – | 133,874 | 5,878 ($T_M$) | 2.53e-06 ($M_{Ch2In}$)<br>2.11e-06 ($M_{In2Ch}$)<br>4.07e-07 ($M_{Ch2In1}$)<br>3.68e-05 ($M_{In2Ch1}$) |
| M3 | *M3_noMig* | 123,988 | 18,776 | 94,639 | 3,729,299 | 2,726,845 | 49,182 | 9,827 ($T_S$) | – |
| | *M3_ancMig* | 94,195 | 13,549 | 93,708 | 3,294,049 | 2,263,451 | 49,890 | 6,905 ($T_S$) | 2.53e-05 ($M_{Ch2In}$)<br>1.31e-05 ($M_{In2Ch}$) |
| | *M3_recMig* | 222,491 | 14,349 | 74,956 | 316,257 | 4,168,176 | 85,373 | 11,564 ($T_S$) | 8.78e-06 ($M_{Ch2In}$)<br>2.30e-05 ($M_{In2Ch}$) |
| | *M3_consMig* | 268,928 | 14,203 | 68,473 | 259,595 | 102,844 | 104,615 | 7,053 ($T_S$) | 3.49e-06 ($M_{Ch2In}$)<br>1.69e-05 ($M_{In2Ch}$) |
| | *M3_difMig* | 164,721 | 11,223 | 73,955 | 314,407 | 88,410 | 93,718 | 4,643 ($T_S$) | 8.91e-06 ($M_{Ch2In}$)<br>1.70e-05 ($M_{In2Ch}$)<br>8.07e-07 ($M_{Ch2In1}$)<br>1.62e-05 ($M_{In2Ch1}$) |
| M4 | *M4_noMig* | 10,207 | 169,827 | 103,378 | 2,060,799 | 55,904 | 35,835 | 415 ($T_{CH}$)<br>1,752 ($T_{IN}$) | – |
| | ***M4_consMig*** | **219,783** | **14,091** | **64,723** | **76,025** | **270,072** | **139,371** | **71,781** ($T_{CH}$)<br>**8,814** ($T_{IN}$) | **6.63e-06** ($M_{Ch2In}$)<br>**1.46e-05** ($M_{In2Ch}$) |
| | *M4_difMig* | 270,855 | 4,581 | 4,185 | 14,035 | 80,690 | 397,667 | 103,262 ($T_{CH}$)<br>1,464 ($T_{IN}$) | 2.27e-06 ($M_{Ch2In}$)<br>4.05e-05 ($M_{In2Ch}$)<br>4.93e-05 ($M_{Ch2In1}$)<br>4.35e-06 ($M_{In2Ch1}$)<br>3.92e-06 ($M_{Ch2In2}$)<br>1.27e-05 ($M_{In2Ch2}$) |
| alternate mutation rate | *M4_consMig** | 97,535 | 5,601 | 21,986 | 24,710 | 45,917 | 80,351 | 31,538 ($T_{CH}$)<br>2,884 ($T_{IN}$) | 1.16e-05 ($M_{Ch2In}$)<br>4.10e-05 ($M_{In2Ch}$) |

**Table S2.** Likelihood estimation for each of 13 demographic models tested with fastsimcoal2, categorized into four groups: *simple split with no size change* (M1), *two-epoch migration models with no size change* (M2), *one size-change models with migration* (M3), and *models with different size-change timing for the Chinese and Indian populations* (M4) (and see Supplementary Figure S1 for detailed model information). For each model, the difference between the maximum observed likelihood and the maximum estimated likelihood (*MaxObsLhood − MaxEstLhood*) is reported. The (*MaxObsLhood − MaxEstLhood*) obtained from the best model, *M4_consMig*, is highlighted in green. Additionally, the best model was rerun using an alternative mutation rate of $1.49 \times 10^{-8}$ per site per generation to account for uncertainty in the mutation rate; this model is referred to as *M4_consMig**.

| model group | model | ***MaxObsLhood - MaxEstLhood*** |
|---|---|---|
| M1 | *M1_noMig* | -120,200.53 |
| | *M1_Mig* | -30,602.60 |
| M2 | *M2_ancMig* | -6,189.39 |
| | *M2_difMig* | -5,180.15 |
| | *M2_recMig* | -31,454.42 |
| M3 | *M3_noMig* | -67,298.69 |
| | *M3_ancMig* | -66,939.64 |
| | *M3_recMig* | -8,172.62 |
| | *M3_consMig* | -3,279.13 |
| | *M3_difMig* | -5,440.94 |
| M4 | *M4_noMig* | -113,859.56 |
| | *M4_consMig* | -2,959.77 |
| | *M4_difMig* | -3,102.08 |
| alternative mutation rate | *M4_consMig** | -1,785.89 |

**Table S3.** Mean $F_{ST}$ values obtained from simulation of the seven best-fitting demographic models inferred from two non-model-based approaches (MSMC2 and Stairway Plot2) and two model-based approaches (fastsimcoal2 [fsc2] and *δaδi*). The $F_{ST}$ value obtained from the best model, *M4_consMig*, is highlighted in green. Additionally, the best model was rerun using an alternative mutation rate of 1.49 × $10^{-8}$ per site per generation to account for uncertainty in the mutation rate; this model is referred to as *M4_consMig**.

| model | Weir and Cockerham's weighted mean $F_{ST}$ |
|---|---|
| empirical data | 0.1476 |
| MSMC2 | 0.0836 |
| Stairway Plot2 | 0.2396 |
| fsc2 - *M1_Mig* | 0.1837 |
| fsc2 - *M2_difMig* | 0.1359 |
| fsc2 - *M3_consMig* | 0.1509 |
| **fsc2 - *M4_consMig*** | **0.1498** |
| *δaδi* - *sym_mig_size* | 0.1435 |
| fsc2 - *M4_consMig** | 0.1493 |

**Table S4.** The estimated best parameter values for each of the best-fitting models across the two non-model-based approaches (MSMC2 and Stairway Plot2) and two model-based approaches (fastsimcoal2 [fsc2] and *δaδi*). Estimated parameters include the ancestral population size ($N_{anc}$), the current Chinese population size ($N_{CH}$), the current Indian population size ($N_{IN}$), the time of the split of the two populations ($T_{DIV}$), and the migration rates $M_{Ch2In}$ (from the Chinese to the Indian population) and $M_{In2Ch}$ (from the Indian to the Chinese population). Parameters not included in a given model are indicated by a "–" symbol. The parameters obtained from the best model, *M4_consMig*, are highlighted in green. Additionally, the best model was rerun using an alternative mutation rate of 1.49 × $10^{-8}$ per site per generation to account for uncertainty in the mutation rate; this model is referred to as *M4_consMig**.

| method | $N_{anc}$ | $N_{CH}$ | $N_{IN}$ | $T_{DIV}$ (in generations) | migration | |
|---|---|---|---|---|---|---|
| | | | | | $M_{Ch2In}$ | $M_{In2Ch}$ |
| MSMC2 | ~120,000 (China)<br>~100,000 (India) | ~30,000 | ~40,000 | ~910 | – | – |
| Stairway Plot2 | ~70,000 | ~120,000 | ~100,000 | ~110,000 | – | – |
| fastsimcoal2 | 64,723 | 219,783 | 14,091 | 139,371 | 6.63e-06 | 1.46e-05 |
| *δaδi* | 84,133 | 80,960 | 7,247 | 58,330 | 2.41e-07 | 2.41e-07 |
| alternative mutation rate | 21,986 | 97,535 | 5,601 | 80,351 | 1.16e-05 | 4.10e-05 |

**1. M1: Simple split with no size change models**

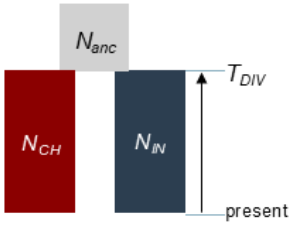


No migration (*M1_noMig*)

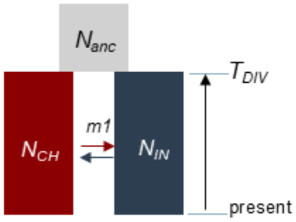


With migration (*M1_Mig*)

**2. M2: Two-epoch migration models with no size change**

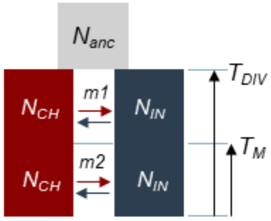


Different migration (*M2_difMig*)

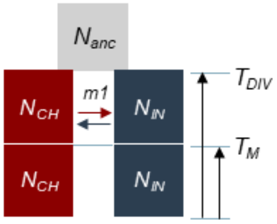


Ancient migration (*M2_antMig*)

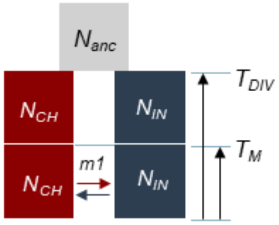


Recent migration *(M2_recMig)*

**3. M3: One size-change models with migration**

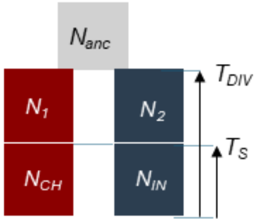


No migration (*M3_noMig*)

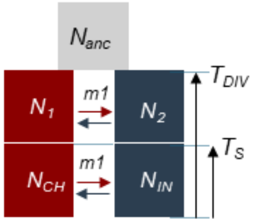


Constant migration (*M3_consMig*)

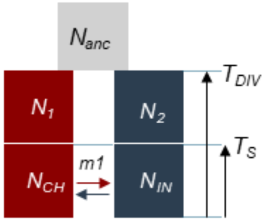


Recent migration *(M3_recMig)*

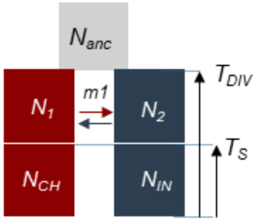


Ancient migration (*M3_antMig*)

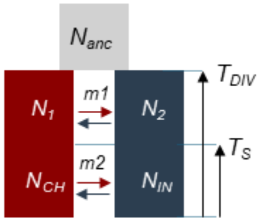


Different migration (*M3_difMig*)

**4. M4: Different time size change models**

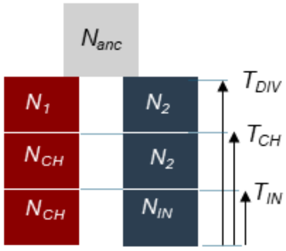


No migration (*M4_noMig*)

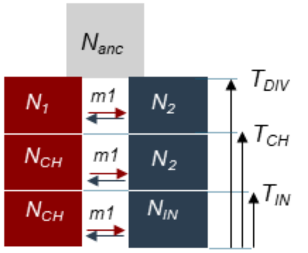


Constant migration *(M4_consMig)*

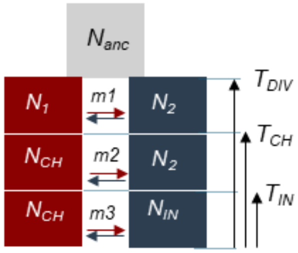


Different migration *(M4_difMig)*

**Figure S1:** Diagram of each of 13 demographic models tested with fastsimcoal2, categorized into four groups: *simple split with no size change* (M1), *two-epoch migration models with no size change* (M2), *one size-change models with migration* (M3), and *models with different size-change timing for the Chinese and Indian populations* (M4). The parameters are the ancestral population size ($N_{anc}$), the current Chinese population size ($N_{CH}$), the current Indian population size ($N_{IN}$), the time of the split of the two populations ($T_{DIV}$), the time of the size change for both populations for M3 models ($T_S$), the time of the size change for the Chinese population in M4 models ($T_{CH}$), the time of the size change for the Indian population in M4 models ($T_{IN}$), the Chinese population size before the size change at $T_S$ or $T_{CH}$ ($N_1$), the Indian population size before the size change at $T_S$ or $T_{IN}$ ($N_2$), the time of migration change for M2 models ($T_M$), and three different migration matrices for different migration time points (*m1*, *m2*, *m3*).

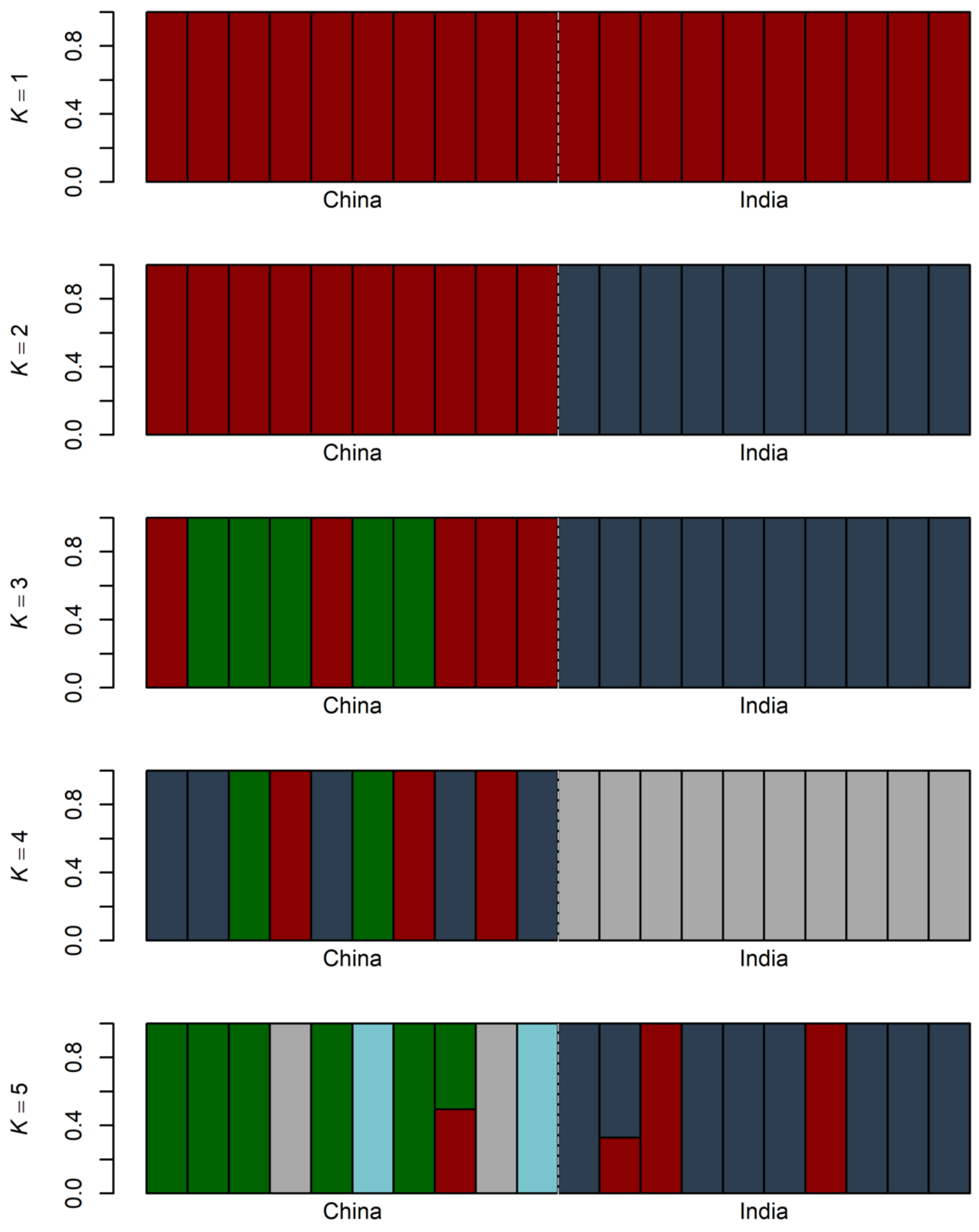


**Figure S2:** Genetic structure of rhesus macaque (*Macaca mulatta*) populations in China and India inferred using ADMIXTURE at $K$ = 1–5. Each vertical bar represents one individual, and colors denote inferred ancestry proportions.

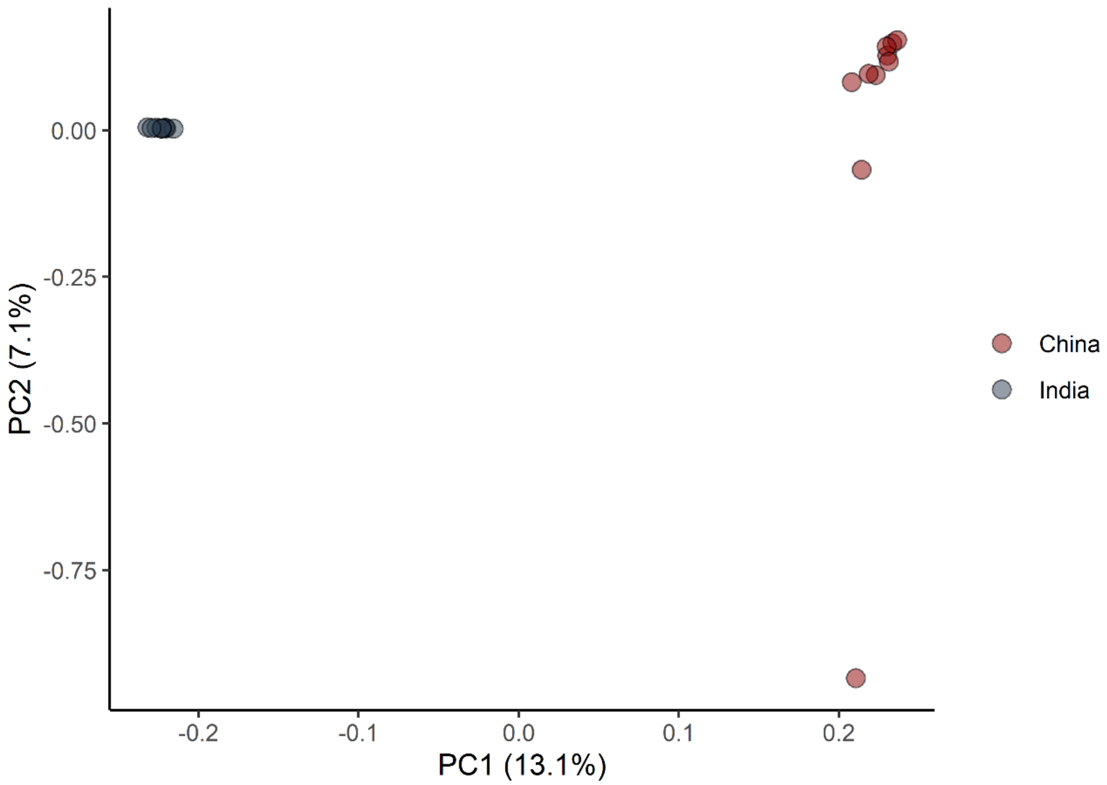


**Figure S3:** Principal component analysis (PCA) of genome-wide neutral variation. Each point represents an individual, and colors indicate population assignment (with the Chinese population shown in red and the Indian population shown in blue). The first two principal components (PC1 and PC2) explain 13.1% and 7.1% of the total genetic variance, respectively. The clustering pattern reflects the genetic differentiation between two populations.

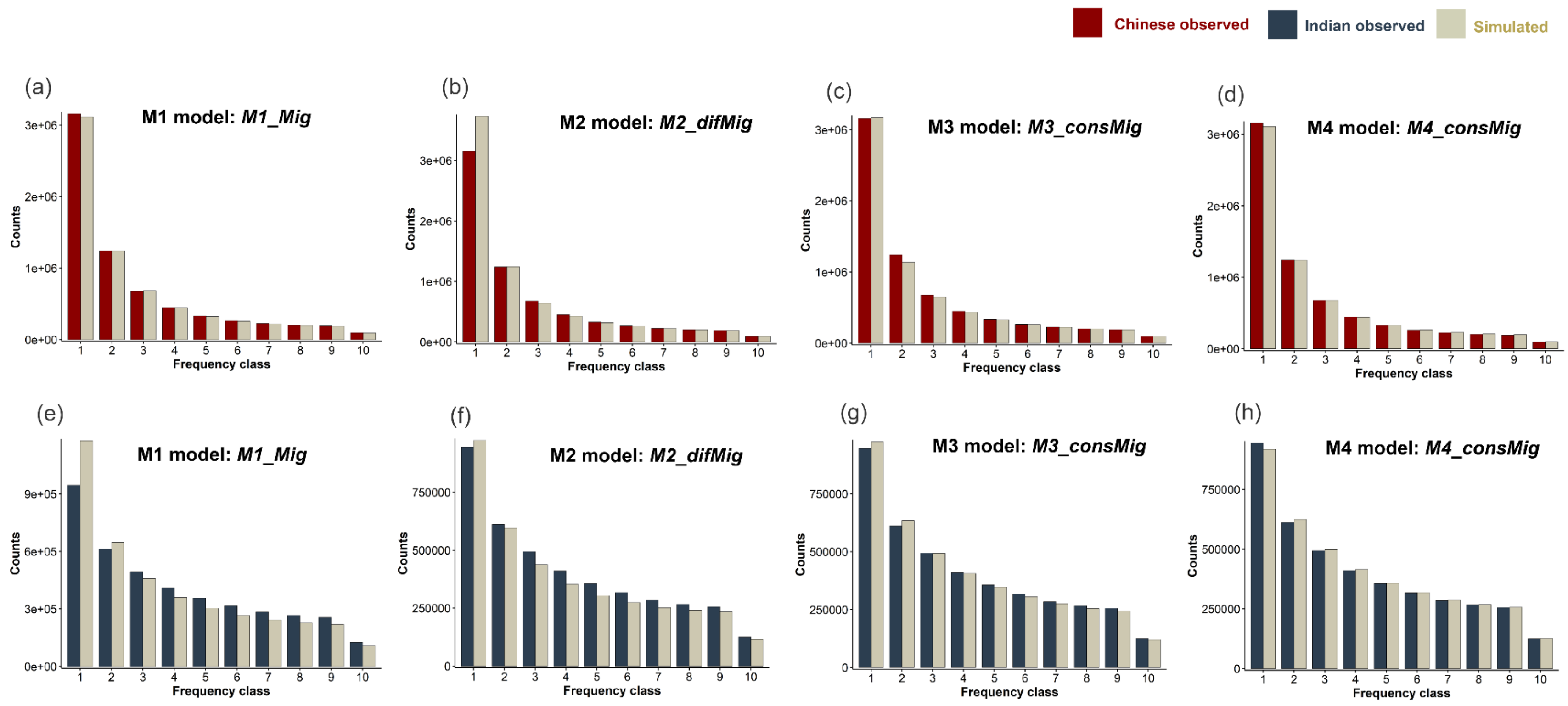


**Figure S4:** The model fit obtained from simulation of the best-fit model from each of four groups inferred with fastsimcoal2 — *simple split with no size change* (M1), *two-epoch migration models with no size change* (M2), *one size-change models with migration* (M3), and *models with different size-change timing for the Chinese and Indian populations* (M4) — compared to the empirically-observed SFS. The top panel (a-d) shows the simulated site frequency spectra (SFS) for the best-fitting models in each group for the Chinese (red) population. The bottom panel (e-f) shows the simulated SFS for the best-fitting models in each group for the Indian (blue) population.

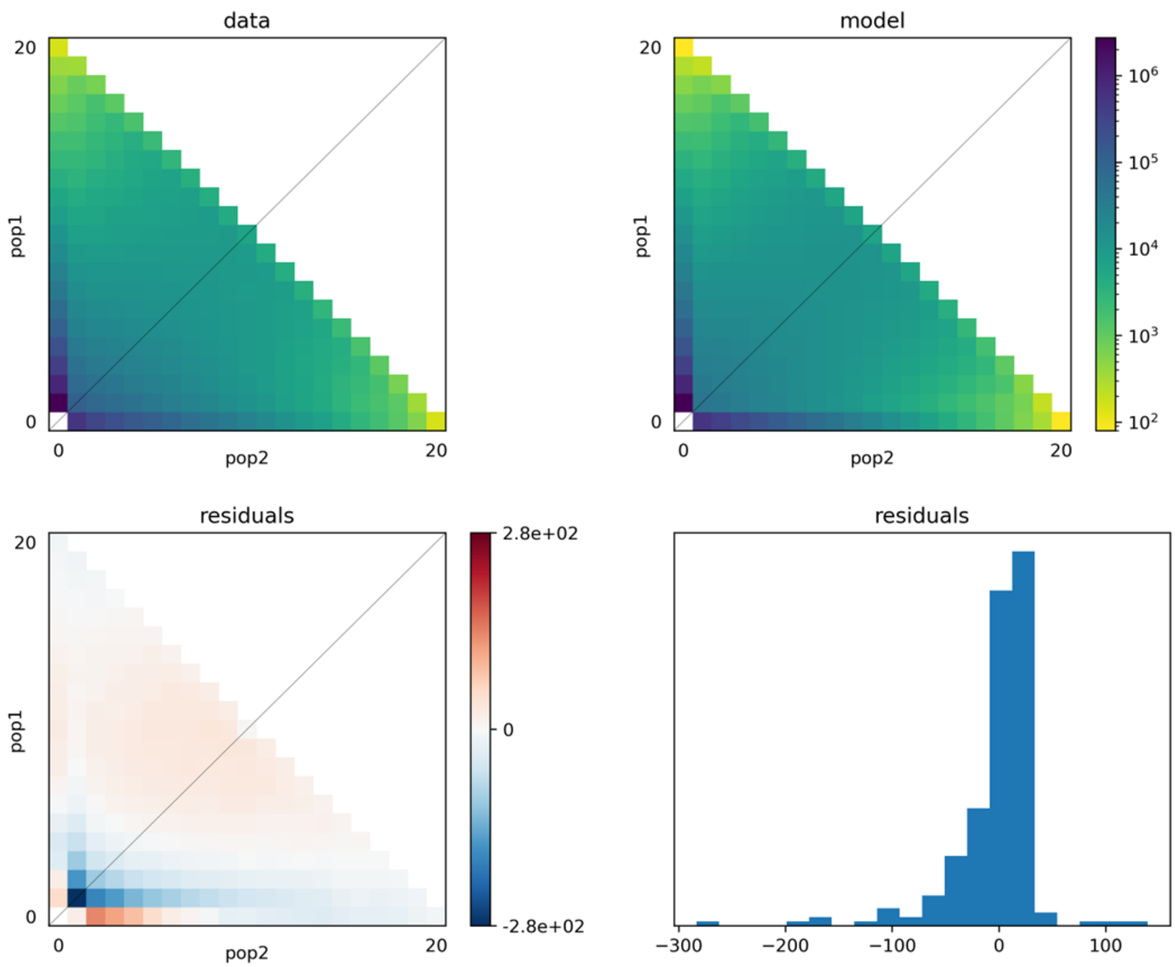


**Figure S5:** Residuals corresponding to the best-fitting *δaδi* model and parameters, with pop1 corresponding to the Chinese population and pop2 corresponding to the Indian population.

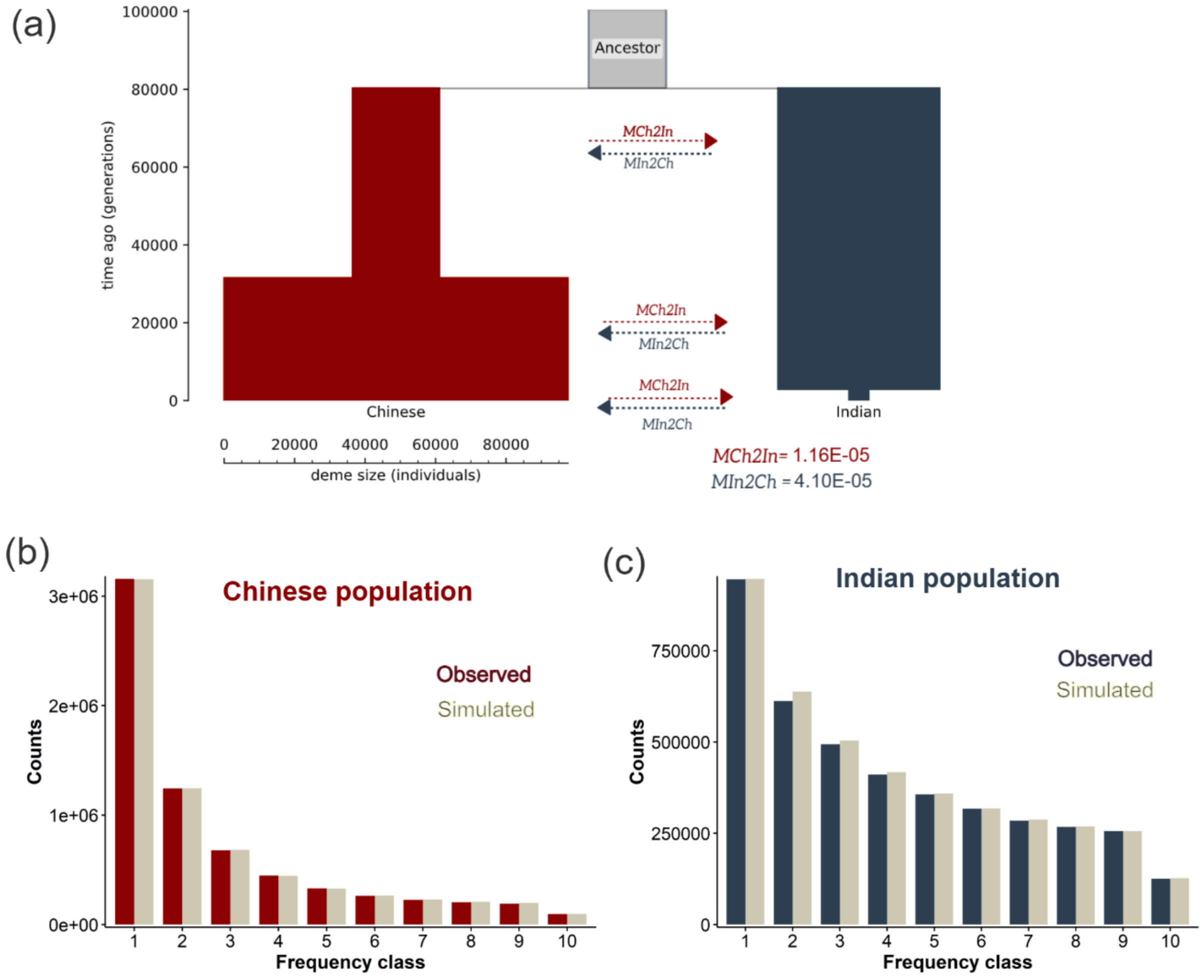


**Figure S6:** (a) Diagram of the best model, *M4_consMig*, rerun with fastsimcoal2 using an alternative mutation rate of $1.49 \times 10^{-8}$ per site per generation to account for uncertainty in the mutation rate. (b-c) Folded SFS for the Chinese (red) and Indian (blue) populations compared to simulations under this model (shown in gray).
